# Skeletonization of Plant Point Cloud Data in Stochastic Optimization Framework

**DOI:** 10.1101/2020.02.15.950519

**Authors:** Ayan Chaudhury, Christophe Godin

**Author notes:** The Authors are with INRIA Grenoble Rhône-Alpes team MOSAIC, Laboratoire Reproduction et Développement des Plantes, Univ Lyon, ENS de Lyon, UCB Lyon 1, CNRS, INRA. Email: *{, }.

## Abstract

Skeleton extraction from 3D plant point cloud data is an essential prior for myriads of phenotyping studies. Although skeleton extraction from 3D shapes have been studied extensively in the computer vision and graphics literature, handling the case of plants is still an open problem. Drawbacks of the existing approaches include the zigzag structure of the skeleton, nonuniform density of skeleton points, lack of points in the areas having complex geometry structure, and most importantly the lack of biological relevance. With the aim to improve existing skeleton structures of state-of-the-art, we propose a stochastic framework which is supported by the biological structure of the original plant (we consider plants without any leaves). Initially we estimate the branching structure of the plant by the notion of β-splines to form a *curve tree* defined as a finite set of curves joined in a tree topology with certain level of smoothness. In the next phase, we force the discrete points in the curve tree to move towards the original point cloud by treating each point in the curve tree as a center of Gaussian, and points in the input cloud data as observations from the Gaussians. The task is to find the correct locations of the Gaussian centroids by maximizing a likelihood. The optimization technique is iterative and is based on the Expectation Maximization (EM) algorithm. The E-step estimates which Gaussian the observed point cloud was sampled from, and the M-step maximizes the negative log-likelihood that the observed points were sampled from the Gaussian Mixture Model (GMM) with respect to the model parameters. We experiment with several real world and synthetic datasets and demonstrate the robustness of the approach over the state-of-the-art.

## I. Introduction

AUTOMATED analysis of phenotyping traits of plants using imaging techniques are becoming off-the-shelf tool for botanical analyses these days ([1], [2]). Although high throughput 2D imaging based phenotyping systems have shown promising results on the accuracy and precision for many cases ([3], [4], [5]), but these systems suffer from the inherent limitations of 2D image based analysis techniques. In recent years, 3D point cloud based analysis is getting extremely popular in phenotyping and agricultural applications [6]. Typical applications of point cloud based phenotyping include plant organ segmentation [7], robotic branch pruning [8], automated growth analysis [9], etc. Many of these applications require skeleton structure of the input point cloud data as a prior for further processing. Also, as a skeleton is a compact representation of the original object, dealing with skeleton requires less computational overhead than dealing with the original point cloud data. That is why skeletons are widely used for varieties of shape analysis tasks in different application areas [10].

In general, a skeleton is a thin structure obtained from an object, which encodes the topology and basic geometry of the object. Ideally, the skeleton should follow the exact centerline of the object. That means within a local neighbourhood, the distance from each skeleton point to the enclosing shape boundaries should be the same. Although skeleton extraction from 2D images is a well studied area, 3D point cloud skeletonization is still an open problem [11]. For the case of plants, skeleton extraction is extremely challenging due to their complex geometry, thin structure, missing data due to self-occlusion, etc. State-of-the-art algorithms ([12], [13], [14]) which are proposed as general skeletonization techniques for regular 3D objects, often produce degraded results when applied to complex plant branching structures. Skeletonization of 3D plant point cloud data is a little worked problem, and only a handful of techniques have been proposed in recent years. These techniques can be broadly classified into two categories: i) initial skeleton construction, and ii) skeleton refinement.

The first type of algorithm refers to skeleton construction from the input point cloud data. A notable work in this category is the method proposed by Xu *et al.* [15]. Initially, a *Riemannian graph* is constructed from the point cloud by connecting the points using nearest neighbour strategy. Then the whole point cloud is clustered based on graph adjacency information and quantized shortest path lengths from the root to all points in the cloud. The center of each cluster is assigned as a skeleton point and the edges are defined as the connection of cluster centers based on their spatial locations. One problem of the technique is that, the skeleton does not maintain the centerdness criteria, and results in a zigzag structure near the junction point where two or more branches meet. Also, the computed skeleton points sometimes end up being outside the boundary of the point cloud (we discuss about these issues in the next section). Bucksch *et al.* [16] proposed a solution of these type of problems by subdividing the point cloud into octree cells and used some local heuristics to form a skeleton graph from the cells. Zhang *et al.* [17] later applied this technique locally to individual branches. However, these methods rely on several heuristic assumptions and require many parameters to be tuned to obtain the desired result. On a different type of approach, *particle flow* based techniques ([18], [19]) are built on the motivation on the process of transport and exchange of energy and water between root, branches and leaves of a plant. However, the skeleton is not guaranteed to follow the actual geometry of the input point cloud data, and thus suffers from biological irrelevance of the results. This technique is mainly used in computer graphics applications. *Space colonization* algorithm ([20], [21]) extracts skeleton from input data by an iterative technique to *eat-up* the points in the cloud starting from the root node at the bottom. The overall technique is a local optimization based strategy. Although the technique can produce visually pleasing skeleton structure, the algorithm might result in creating branching structure in the wrong directions, and there is no mechanism to perform backtracking to correct the biological irrelevance of the branches.

The second type of approach of skeletonization is based on the motivation to improve the initial skeleton. Livny *et al.* [22] proposed a series of optimization techniques to smooth the branches of a skeleton in a realistic manner. However, the optimization strategy does not involve prior botanical knowledge of plants. Branch smoothing is performed independently without taking into consideration the original data, and can produce over-smoothing or under-smoothing of branches which suffer from botanical inconsistency of the results. In a similar line of work, Wang *et al.* [23] proposed a strategy to refine an existing skeleton to handle the case of occlusion and missing data. The approach is based on a combined local and global optimization strategy, where the coarse skeleton is pushed to move towards the original point cloud, and the original point cloud is forced to contract towards the skeleton points. The method is explicitly built to handle the occlusion cases and overlooks factors like zigzag problem and biological irrelevance of the skeleton. In a recent work, Wu *et al.* [24] proposed a skeleton refinement technique for Maize plants. The issue of centerdness criteria and zigzag problem is explicitly addressed. The refinement technique is based on a local neighbourhood based approximation strategy and some heuristics are used. The method is tested only on a single plant species. Other types of recent skeletonization techniques are shown to be successful for some applications [25], [26], where it is assumed that the point cloud is voxelized along with voxel connectivity information, instead of considering raw point cloud data without any connectivity information.

We propose a skeletonization technique that belongs to the second category. The aim is to improve an existing skeleton obtained from the state-of-the-art by maintaining the biological relevance with the input point cloud data. Initially we represent a coarse skeleton as a tree structure which consists of superposition of parametric curves having uniform density of discrete points. Then we propose a stochastic optimization technique to transform the skeleton points so that the transformed points maintain the centerdness criteria as well as the biological relevance with the input data. Existing skeleton refinement techniques typically aim at improving an initial coarse skeleton by optimizing an objective function. Often the objective function is constructed using biological prior, and the optimization does not make use of the original input point cloud data ([22]). Other approaches ([23]) deform the original point cloud data during the optimization process. However, these types of approaches do not guarantee the optimized skeleton to be geometrically consistent with the original point cloud data. One of the main strength of our approach is the exploitation of the input point cloud data during the optimization process. This explicitly helps the skeleton to retain biologically supported structure.

## II. Problem Statement

The motivation of this work is to extract the skeleton from plant point cloud data. Before we discuss our model in the next section, we first demonstrate the limitations of the existing approaches on real datasets. We identify the following 3 problems associated with the existing techniques, as shown visually in Figure 1.

**Fig. 1.**
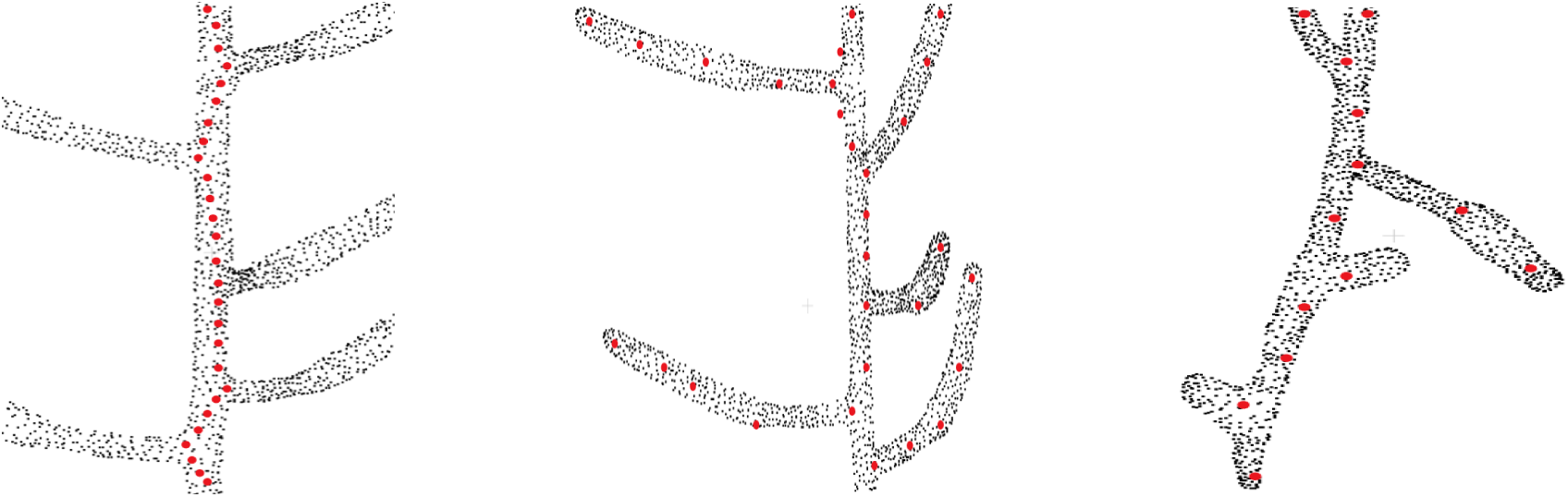
Problems with the existing approaches on skeletonizing plant point cloud data. Original point cloud is shown as black point cloud and the skeleton points are shown as red points. Left: The problem of zigzag structure, where the skeleton does not follow the centerline of the stem and tends to deviate towards the branching point [15] (only the main stem skeleton is shown in the figure). Middle: The problem of biologically irrelevant skeleton points which falls beyond the boundary of the input data (shown at the top part), and inability to capture the geometric details for some branches [27], [7]. Right: Overlooking tiny geometrical structures [15].

### A. *The* Zigzag *Structure Problem*

One of the typical problems associated with skeletonizing plant point cloud data is the deviation of skeleton points from the centerline of the branch towards the junction point where two or more branches are joined [15], thus creating a zigzag type skeleton as shown in the left of Figure 1. The zigzag problem occurs mainly because of the local nature of the algorithm. During the clustering phase, the data is quantized into several levels, and the center of each cluster is computed as a skeleton point. For a simple branch, the cluster center is computed at the center of the branch. However for the case of a bifurcation, the cluster center tends to shift towards the bifurcating branch, thus resulting in a zigzag like structure. Ideally the skeleton points should follow the exact centerline of the branch representing the actual geometry of the plant. However, the zigzag skeleton structure is a wrong approximation of the plant geometry, which results in wrong phenotyping parameter estimation in quantitative measurement of phenotypes such as internode distance, branch length, etc. Although this problem is handled locally in [24], the method is based on many heuristics to skeletonize a handful of Maize plant point cloud data.

### B. Invalid and Inaccurate Geometry Estimation Problem

As shown in the middle of Figure 1, sometimes skeleton points do not lie within the enclosing boundary of the original point cloud data [27], [7]. This results in invalid geometry estimation of the input data. For example, in the upper part of the figure we can see the red dots which are located outside the boundary of the point cloud. Also for the case of curved branches, insufficient number of skeleton points are not able to represent the actual geometry of the branch. This is shown at the bottom of the figure where the curved branches are not represented by the skeleton points. One way to handle this problem might be to generate more skeleton points in the curved areas using interpolation or similar point approximation strategies. However, the generated points will be independent of the original point cloud data, and will fail to reconstruct the original geometry of the branch.

### C. Inability to Handle Tiny Structures

Tiny branches are often treated as noise in the point cloud data, and the extracted skeleton fails to represent these branches. This is shown at the right of Figure 1. Two tiny branches are completely overlooked in terms of skeleton representation of the input data [15].

## III. Building and Optimizing the Skeleton

### A. Skeleton Intialization

Initially we obtain a coarse skeleton graph of the input point cloud 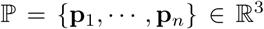 by the method of Xu *et al.* [15]. Note that other skeletonization algorithms can also be used to obtain the coarse skeleton, but we have used this method because the algorithm is simple, fast, works reasonably well, and the open source implementation is available^1^. Next we remove the cycles in the graph by the method of Yan *et al.* [28]. The resultant skeleton point set is a rooted tree graph embedded in ℝ^3^. In this work, we represent the input skeleton and its branching structure as an *axial tree* ([29], [30]).

Formally, let 𝓖 = (*V, E*) be a graph where *V* = *v*_*i*_}_1≤*i*≤*I*_ (where I is an integer ≥ 1) is the set of vertices, and *E* = (*v*_*i*_, *v*_*j*_) is the set of edges connecting to the ordered pairs of vertices *v*_*i*_ and *v*_*j*_. The vertex *v*_*j*_ is the child of *v*_*i*_ denoted as, child(*v*_*i*_) = {*v*_*j*_ ∈ *V*|(*v*_*i*_, *v*_*j*_) ∈ *E*}, and the vertex *v*_*i*_ is the parent of *v*_*j*_ denoted as *v*_*i*_ = parent(*v*_*j*_). A rooted tree graph is a graph that contains no cycle, only one connected component, and there exists a unique vertex in *V*, called the *root* that has no parent. The *descendants* of a vertex *v*_*i*_ ∈ *V* is defined as the set of vertices that belong to the branching system starting from *v*_*i*_ excluding itself.

#### Definition 1

Axial Tree, adapted from [30]: An *Axial Tree* 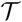 is defined as a tree graph along with a *successor* function *succ* associated with each vertex in the graph defined as,

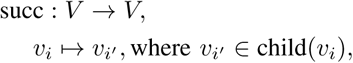

where *v*_*i′*_ is called the successor of *v*_*i*_.

From the definition, we see that all the vertices *v*_*i*_ ∈ *V* in an axial tree, there exists at most one successor *v*_*i′*_ ∈ *V*. Note that there may be vertices that do not have a successor in *V* (such as leaf nodes in particular). Maximal set of vertices connected by successor relationship are paths in 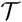 (called the *axes* of 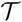). For example in Figure 2(a), the axes are denoted as *a*1,…, *a*9. In this paper, a skeleton is thus represented by an axial tree 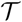 for which the vertices correspond to the points in the skeleton, and the edges as the ordering of these points within their axis. The 3D coordinates **s**_*i*_ of each point is attached to the corresponding vertex *v*_*i*_.

**Fig. 2.**
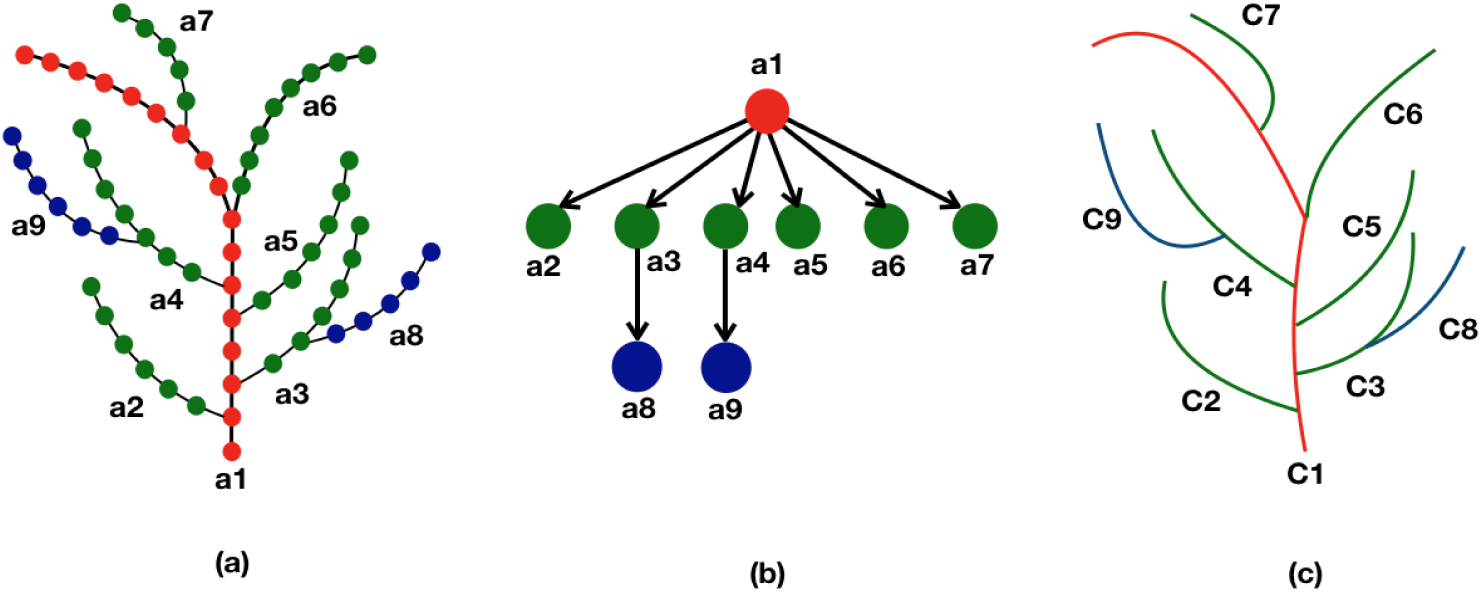
Representation of skeleton tree graph as a curve tree via quotient graph. Left: each branch (*a*1,…, *a*9) in the skeleton tree represents an axis where the discrete points are shown as coloured dots using a hierarchical colour convention. The main stem is represented as red dots, next level branches as green dots, and further level branches as blue dots. The successor of a node of certain colour is coloured with the same colour. Middle: each node in the quotient graph represents each branch (or axis) of the skeleton graph, and the directed edges represent their hierarchical relationship in the skeleton structure. Right: corresponding curve tree where each branch in the skeleton graph is approximated as a parametric curve (*C*1, …, *C*9).

In a given axis *a* = {*v*_1_,…, *v*_*k*_} (with slight abuse of notation), we define the *bearing vertex w* as, *w* = *b*(*a*) ⇔ *w* = parent(*v*_1_), if it exists. The *order* relationship between the vertices derived from their order on the path in the axis is defined as, *v*_*i*_ < *v*_*j*_ ⇔ *v*_*i*_ is an antecedent of *v*_*j*_.

### B. Parametric Representation of the Tree Skeleton

Now we want to define for each axis in the axial tree a parametric model represnting its geometry. For this, we first define a conceptual model representing the branching geometry of the skeleton, and then explain how it is constructed from the data. Let us first introduce the following definitions.

In an axial tree 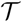, we say that two vertices *v*_*i*_ and *v*_*j*_ are in *equivalence relationship* (denoted as *v*_*i*_ ≡ *v*_*j*_) if and only if *v*_*i*_ and *v*_*j*_ belong to the same axis of an axial tree. This allows us to define the notion of quotient tree graph as follows.

#### Definition 2

Quotient Tree Graph, adapted from [30]: Given an axial tree graph 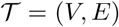, a *Quotient Tree Graph* 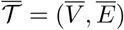 is the quotient graph of the axial tree graph 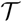, which is made of the axes of 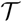, and is also a tree graph.

The vertices in an axial tree having equivalence relationship with each other, are collapsed to a single node in the quotient graph (Figure 2(b)).

Now consider a family of curves 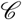 defined in ℝ^3^. With each vertex 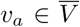 in the quotient graph 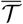, we associate a curve 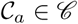 defined as,

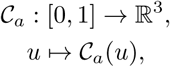

where *u* ∈ [0, 1] is the parameter of the curve. Curves on the axes must respect the following attachment conditions to one another. For any two vertices *a*_*i*_, 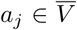 such that *a*_*j*_ = parent(*a*_*i*_), then the following condition holds:

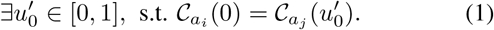

Ideally, we want to represent the skeleton as a union of curves, one curve is being attached with a vertex of the quotient tree graph, where the order of the branching points in the skeleton is the order of vertices in the quotient graph for the attachment of the curves. In order to achieve this, the following condition must hold true in 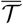. Consider the axes *a*_*i*_, *a*_*j*_, *a*_*k*_, where *a*_*i*_ = parent(*a*_*j*_) and *a*_*i*_ = parent(*a*_*k*_). Let *w*_*j*_ and *w*_*k*_ be the respective bearing vertices for the axes *a*_*j*_ and *a*_*k*_. Then the following condition holds true:

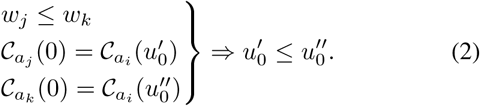

#### Definition 3

Curve Tree: A *curve tree* 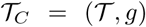 is an axial tree 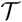 augmented with a mapping *g* that attaches one parametric curve to each vertex of 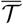, and verifying the attachment conditions in Equation 1 and 2 on its quotient tree graph. The mapping *g* is defined as,

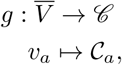

where the curve 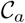 is associated with each vertex 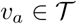. In order to instantiate the curve associated with each axis *a*, we choose a family of curves that may be parameterized using the 3D points {*v*_1_, …, *v*_*k*_} of the axis *a*.

### C. Estimation of the Curve Tree Parameters

From a given axial tree, we can define a curve tree by attaching a parametric curve to each axis *a* of the quotient tree graph from a chosen family of parametric curves 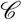.

In this work, we choose to model curves representing axes using the family of splines. Splines are powerful mathematical tools to approximate curves and surfaces. In the general case, a spline curve 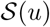 defined by (*n* + 1) control points^2^ *P*_0_, *P*_1_, …, *P*_*n*_ can be defined as,

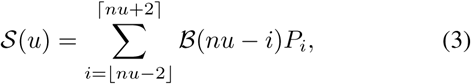

where 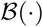 is the blending function of the spline, and *u* ∈ [0, 1] [31]. For a cubic B-spline, at most four nearest control points are used to compute the blending function for one spline point. We aim at resampling each branch of the curve tree, where the number of points in a branch can be modelled to the desired number. We have used the following strategy. Let the Euclidean distance (we are not taking into account the branch bending in the distance computation) between the endpoints of a branch is *d* (in mm). Then we approximate the number of spline points as *d/d*_0_, where we use *d*_0_ = 1.0. Depending on the application, higher or lower density of points can be achieved easily by changing the value of *d*_0_ to a lower or higher value, respectively. Number of spline points can be greater than, equal to or less than the number of control points. Any curve thus can be represented as the polyline joining continuous set of spline points. Although cubic B-splines are widely used in modelling curves, we model the branch curves as *β*-splines, which provide an intermediate representation between approximation and interpolation and thus captures the local geometry better than cubic B-spline. Given the fact that *β* splines are 1^*st*^ and 2^*nd*^ order continuous [32], we can approximate the branches as smooth enough. The blending function of a *β*-spline denoted as *β*(*v*), parameterized by *v* is given by,

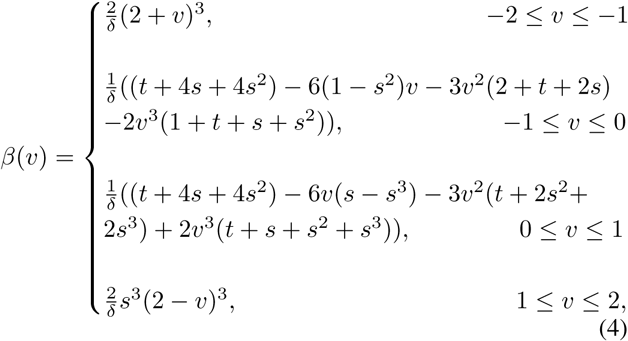

where *v* = *nu* − *i*, *t* is the tension parameter and *s* is the skew parameter ([32], [33]), which are kept constant. We used *t* = 10 and *s* = 1 for all of our experiments. While the skew parameter is always set to 1 to ensure the smoothness of the curve, we have tested different values of *t* on several datasets. From empirical observation, we notice that keeping *t* ∈ [5, 15] produces best approximation of the curve (very low error value of mean square distance between the original points and generated points), while other values of *t* outside this range yields high error value (the error corresponding to *t* = 10 was the minimum). *δ* is given by *δ* = *t*+2*s*^3^+4*s*^2^+4*s*+2. Basically the blending function is defined for 4 discrete intervals (2 in each side) around each control point. While computing the spline points near the endpoints, we use the trick of truncating the generated spline points to the endpoints of the curve in order to prevent the generated points to go beyond the boundary of the curve.

At this stage, the curve tree represents a coarse skeleton of the plant having uniform distribution of discrete points. However, the skeleton still has the zigzag structure and non-centerdness problem. We introduce a stochastic optimization approach to improve the current skeleton to follow the geometry of the original point cloud in order to obtain a biologically relevant skeleton of the input point cloud data.

### D. Stochastic Modelling

Ideally, the skeleton points should follow the exact centerline of the input point cloud data. For the case where the branches of the plant are cylindrical, the skeleton should follow the central axis of the cylinder in each branch. In case of cross-sectional branches, the skeleton should go through the middle of the points by maintaining equidistance criteria from the enclosing boundaries. The initial skeleton suffers from the problem of not maintaining the centerline criteria as defined above, thus resulting in having points which are deviated from the centerline. Our goal is to improve the deviated skeleton by moving the points towards the centerline and maintain the geometry of the branching structure of the plant using a stochastic approach.

We frame the problem of skeleton refinement as a transformation estimation problem. We aim at moving (or transforming) the discrete points of the skeleton towards the original input point cloud data, so that the skeleton points get aligned with the centerline of the original point cloud data. We consider points in 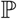 (defined in Sec. III) as the observations of a Gaussian Mixture Model (GMM), and the centroids of the Gaussians are considered as the skeleton point set. Now in order to estimate the correct location of the centroids, we aim at finding which Gaussians the points in the skeleton point set are sampled from. Initially, the discrete points of the resampled curve tree represents the initial location of the Gaussian centroids. Then we iteratively use the Expectation Maximization (EM) algorithm to automatically approximate the optimal centroid locations and reparameterize the Gaussians in each iteration by maximizing a likelihood function. However, the problem is highly non-rigid in nature. Global transformation of points is not sufficient to achieve the optimal solution, and we aim at estimating transformation for every point in the skeleton. Exploiting the recent advancements of Gaussian Mixture Model based approaches ([34], [35], [36]), we formulate the problem in a probabilistic framework.

We denote the original point cloud 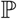 as the *fixed* point set, and the skeleton point cloud 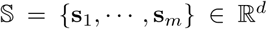 of the resampled curve tree as the *moving* point set, where *d* is the dimension of the point cloud (which is 3 in our case). In a typical case of point set registration, *m* ≈ *n*, and there is a correspondence between most of the point pairs (**s**_*i*_, **p**_*j*_). However in our case *m* ≪ *n*, and instead of one-to-one correspondence, a group of points in the original point set corresponds to a single point in the skeleton point set. Let **Θ** be the set of unknown model parameters (we define **Θ** later). We estimate the centroid locations 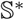 from the optimal set of model parameters **Θ**∗, which is obtained by minimizing the following negative log-likelihood,

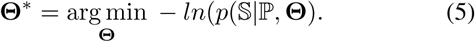

This type of problem can be solved by the classical framework, where the E-step estimates which Gaussian the observed point cloud was sampled from, and the M-step maximizes the negative log-likelihood that the observed points were sampled from the GMM with respect to the model parameters. Each **s**_*i*_ is assumed to be the centroid of a Gaussian, and the corresponding **p**_*j*_’s are assumed to be normally distributed around **s**_*i*_. The probability density function is then given by,

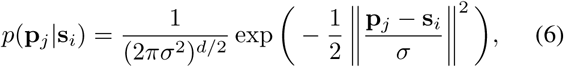

where *σ* is the covariance of the Gaussian, whose initial value *σ*_(0)_ is set as,

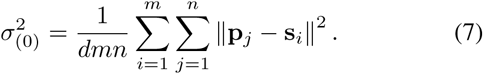

Now we introduce outliers in the model, assuming the outlier probability to have uniform distribution for all data points. Let *ϵ* ∈ [0, 1] be the weight of the outlier distribution in the original point set (that means, each point has the probability *ϵ/n*), and (1 − *ϵ*) be the weight in the skeleton point set. Then the mixture model takes the form,

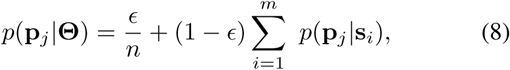

where **Θ** = {*σ*^2^, *ϵ*} is the set of model parameters, which we wish to estimate in EM framework in order to minimize the negative log-likelihood energy of Equation 5. Next, we propose a membership probability 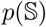 for the GMM components. Instead of assigning equal probability for all the points, a weight factor is used to indicate the probability of point correspondence between the two point sets. The probability that the point **p**_*j*_ is an observation of the Gaussian of point **s**_*i*_ is defined by *α*_*ij*_ as the membership probability. We use a local Principal Component Analysis (PCA) based technique to estimate the membership probability as follows. Considering a local neighborhood 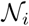 around the *i*-th point defined as the points within a ball of radious *r*, the 3 × 3 covariance matrix 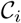 is computed as,

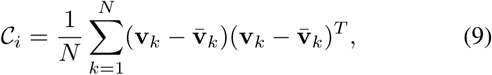

where *v*_*k*_ are the points in the local neighbourhood of *i*-th point, N is the number of points in the neighbourhood, and 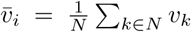. If the eigenvalues of the matrix 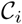 are λ_0_, λ_1_, λ_2_ (where λ_0_ ≤ λ_1_ ≤ λ_2_), then we compute the normalized eigenvalueof the neighbourhood as,

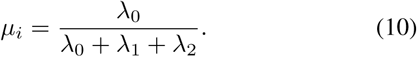

Note that instead of normalizing the lowest eigenvalue, other eigenvalues can also be normalized to compute *µ*_*i*_. We use *µ*_*i*_ as an estimate of the local structure of the neighbourhood of *i*-th point. Similarly we compute *µ*_*j*_ for the other point set. Large difference of *µ* = |*µ*_*i*_ − *µ*_*j*_| between the local neighbourhood is penalized by assigning low probability, whereas small difference contributes to high probability. The membership probability that *i*-th point corresponds to *j*-th point is computed as an exponentially decaying function as,

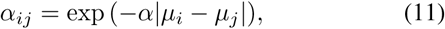

where *α* is a tuning parameter (set to 1 for all of our experiments).

Along with the membership probability in the negative log-likelihood, we minimize the following energy by EM algorithm,

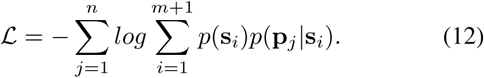

#### E-step

In the E-step, the current parameter values (the “*old”* set of parameters) are used to estimate the posterior distribution *p*^old^(**p**_*j*_|**q**_*i*_) using Bayes’ theorem as,

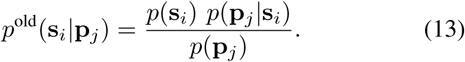

In order to evaluate the probability *p*^old^ in the above equation, we use the expressions from Equation 6, 11, and 8. The first term in the numerator is the probability of correspondence of **p**_*j*_ with **s**_*i*_. This term is estimated as *α*_*ij*_ as in Equation 11. Incorporating the outliers *ϵ* in the model, the second term in the numerator is estimated as (1 − *ϵ*)*p*(**p**_*j*_|**s**_*i*_). The denominator is obtained from Equation 8. Hence the expression for *p*^old^ is written as,

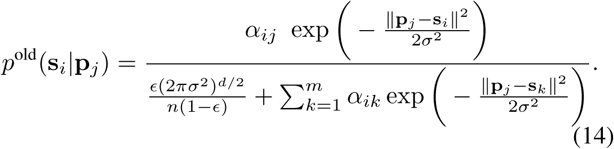

#### M-step

In the M-step, “*new”* parameter set is obtained by minimizing the expectation of the complete negative log-likelihood function of Equation 12 as,

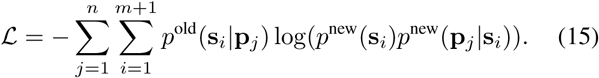

Ignoring the constants independent of **Θ**, we can write the expression for complete likelihood 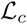 as [36],

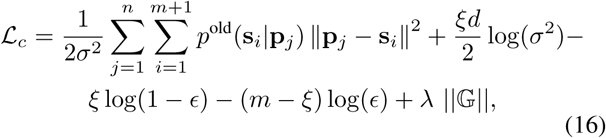

where 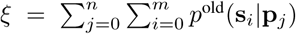, and the last term is a regularizer weighted by a factor *λ* to restrict the spread of the data by the transformation. For a skeleton point **s**_*i*_ and the corresponding transformed point **s′**_*i*_ is regularized as the Gaussian kernel, 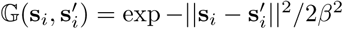, where the variable *β* controls the spread of the data. If the value of *β* is too small, the transformation will have negligible effect on the data, while large value of *β* will result in badly scattered points. Minimizing 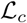 decreases the negative log-likelihood of the energy function.

The EM algorithm iterates until there is not sufficient update of the parameters (we keep the tolerance at 10−^4^). The updated location of skeleton point set S is the optimal transformed point set 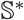.

## IV. Experiments and Analyses

We performed several experiments with real world and synthetic datasets in both qualitative and quantitative manners. More specifically, we have experimented with the data from 3 datasets. The first dataset we have used is the PlantScan3D dataset [37], which contains different varieties of plants including Cherry, and Apple tree point cloud data. Next, we used the dataset from Tabb *et al.* [26]. The dataset contains ‘Fuji’ apple tree data in voxel format. We have extracted the raw point cloud from the data, and performed experiments without using any connectivity information from the voxel structure. Finally, we have generated point cloud data by scanning real Arabidopsis thaliana plants. For all the datasets, we obtained the initial coarse skeleton using the method of Xu *et al.* [15], and performed the proposed optimization technique. Although other types of skeletonization methods can be used to obtain the initial skeleton, the result of the optimization algorithm depends on the quality of the initial skeleton. We have performed the optimization by using Space Colonization algorithm ([20], [21]) as the initial skeleton, where we selected the set of parameters which produced best results. We noticed that in many cases, the results suffer from problems where the skeleton points are too far from the centerline. Especially the branch junction points are located out of the boundary of the point cloud in many cases. The optimization performs poorly in these cases than using the method of Xu *et al.* [15] as the initial skeleton. We used Python 2.7 version along with PlantScan3D library for the implementation. The experiments were performed in a MacBook Pro with 2.2 GHz Intel Core i7 processor and 32GB DDR4 RAM. The datasets in Figure 4 contain about 124k, 30k, and 18k points in the Apple tree, Cherry tree and Arabidopsis data, respectively. The respective computation times are 68 minutes, 81 seconds, and 34 seconds.

First we demonstrate some qualitative results in Figure 3, where we show the skeletonization results on Arabidopsis plant and Cherry tree point cloud data. For visual clarity, we show part of the original data for the Arabidopsis plant. Similarly for the Cherry tree data, we removed the noise and truncated part of a branch to demonstrate the quality of the skeleton. In Figure 4, we show the full skeleton structures along with the original point cloud data for a ‘Fuji’ apple tree, cherry tree, and arabidopsis plant, respectively.

**Fig. 3.**
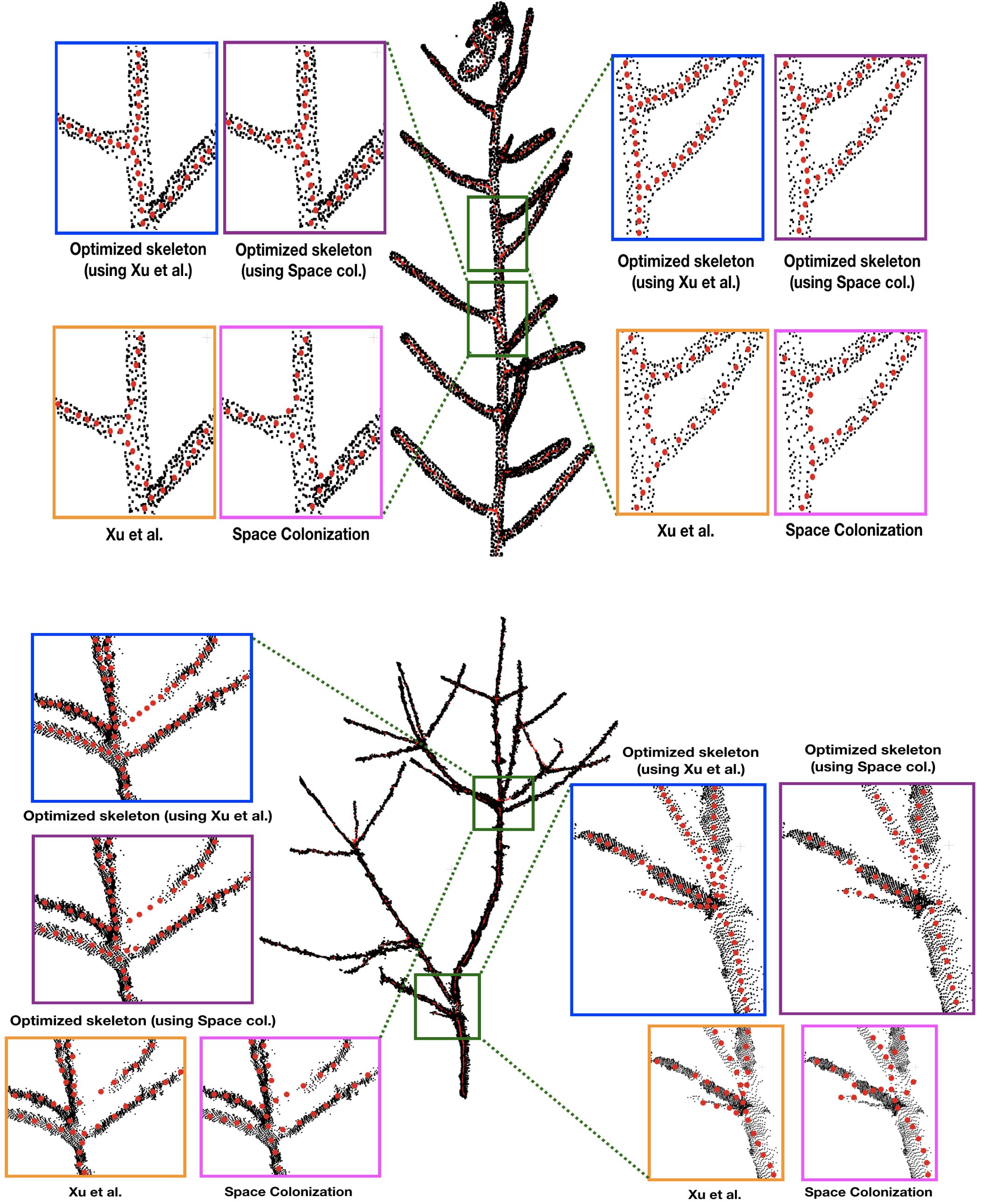
Skeletonization results on 2 datasets: Arabidopsis plant (top) and Cherry tree (bottom) point cloud data. Note that the red skeleton points lie inside the plant stem, and that is why the visibility is not always clear due to the occlusion by the original (black) point cloud. We show some parts of the data by zooming-in and showing the skeleton points at the surface of the plant by taking the points out of the plant.

**Fig. 4.**
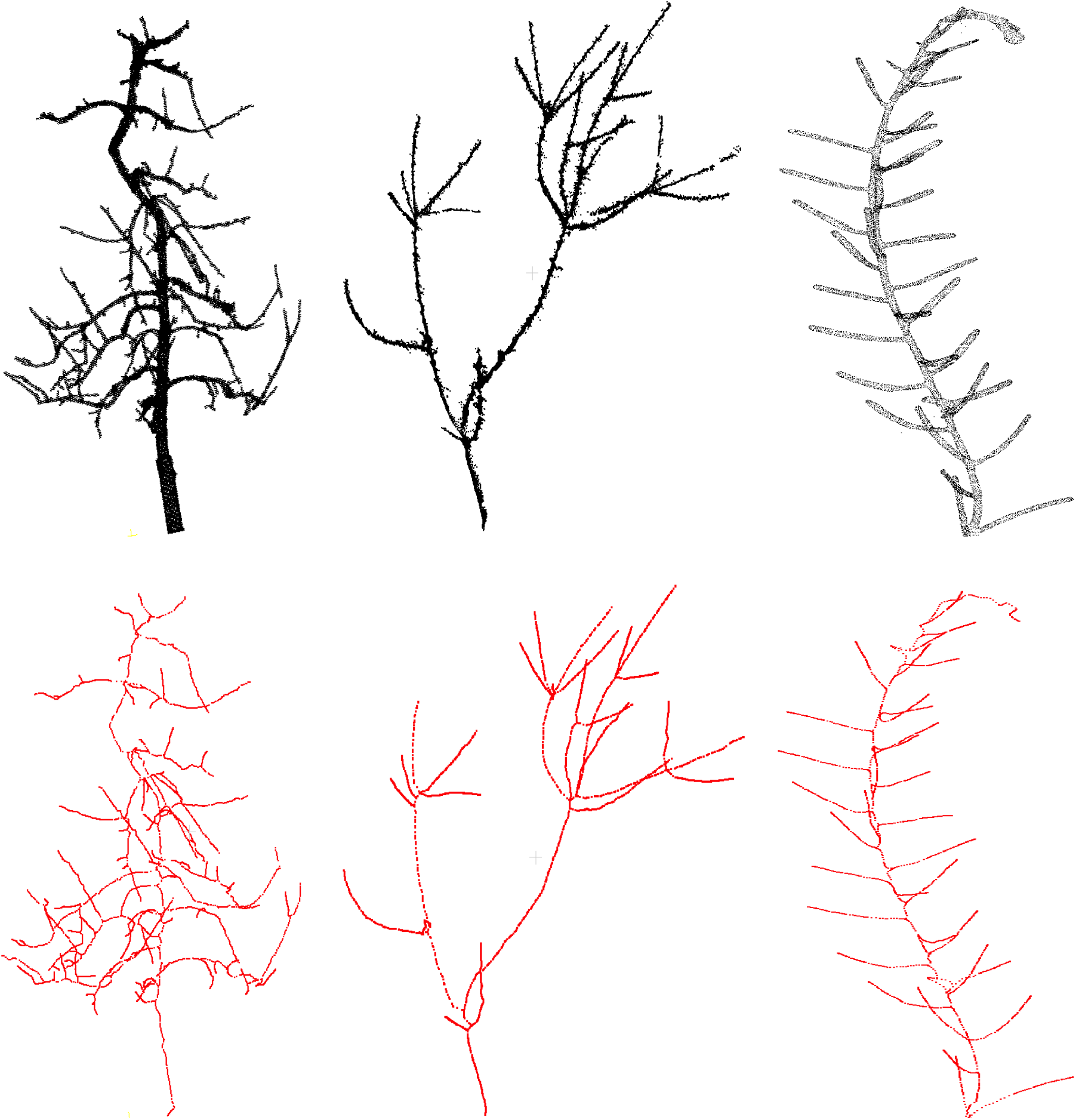
Results of skeletonization on the whole plant point cloud data. Left: dataset from Tabb *et al.* [26], middle: apple tree dataset from INRIA Plantscan3d dataset, right: arabidopsis data.

Next, we demonstrate the quantitative results of different plant phenotypes obtained from the computed skeleton. We have generated ground truth of junction points of the branches (branching point) for 3 cases: synthetic data, Arabidopsis plant data, and Cherry tree data. We notice that the quality of the skeleton explicitly affects the location of the branching point and the relative length of branch segment between the branching points with respect to the average branch length. In order to evaluate the quality of the skeleton, we use two metrics based on the above mentioned factors to quantify the errors. The first metric is the deviation of the computed junction point from the ground truth location, and the second metric is the difference between the ground truth branch segment length and the computed branch segment length with respect to the average branch length. We compare our results with the state-of-the-art skeletonization algorithms which are built specifically for plants, consider raw point cloud data as input to the algorithm, and the implementation is publicly available. More specifically, we compare our algorithm with Xu *et al.* [15] and Space Colonization algorithm [20], [21] implemented in PlantScan3D library. In Figure 5, we show the error bar plots for relative distance error of junction point location and branch segment length error in top and bottom row respectively.

**Fig. 5.**
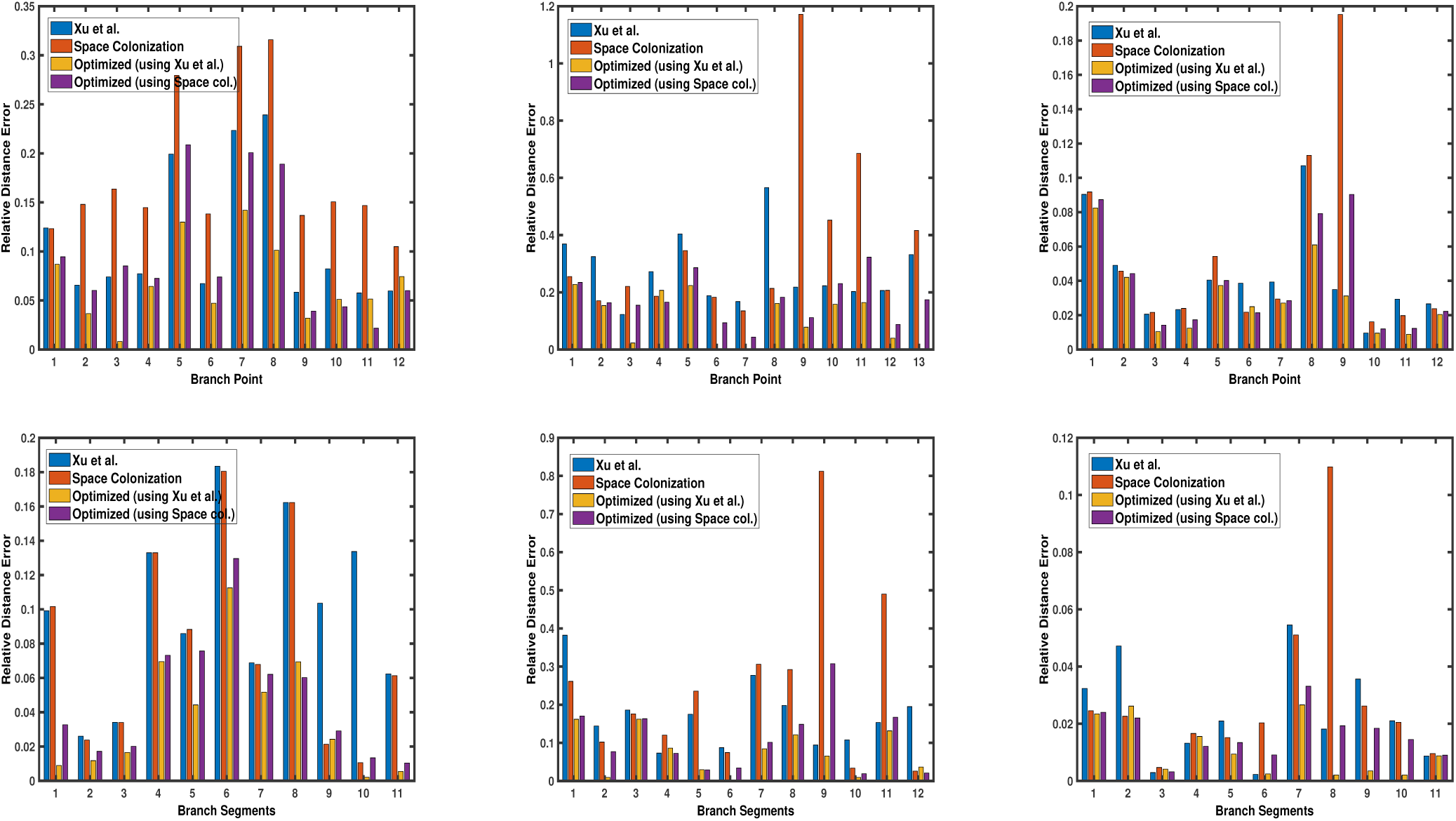
Error bar graphs for relative junction point location (top row) and branch segment length (bottom row) for 3 datasets: a synthetic data, Arabidopsis plant data (shown in each column), and Cherry tree data. In each graph, the vertical bar represents the value of error with respect to the groundtruth. Performance of each algorithm is shown in different colours, as defined by the legend in each graph.

We also show the statistical distribution of the corresponding error values in the box plots of Figure 6. Similar to Figure 5, the top row shows the results of junction point location errors and the bottom row shows the results of relative branch segment length errors. As can be seen from the figure, the span of error values for the proposed optimization algorithm is always less than the other methods.

**Fig. 6.**
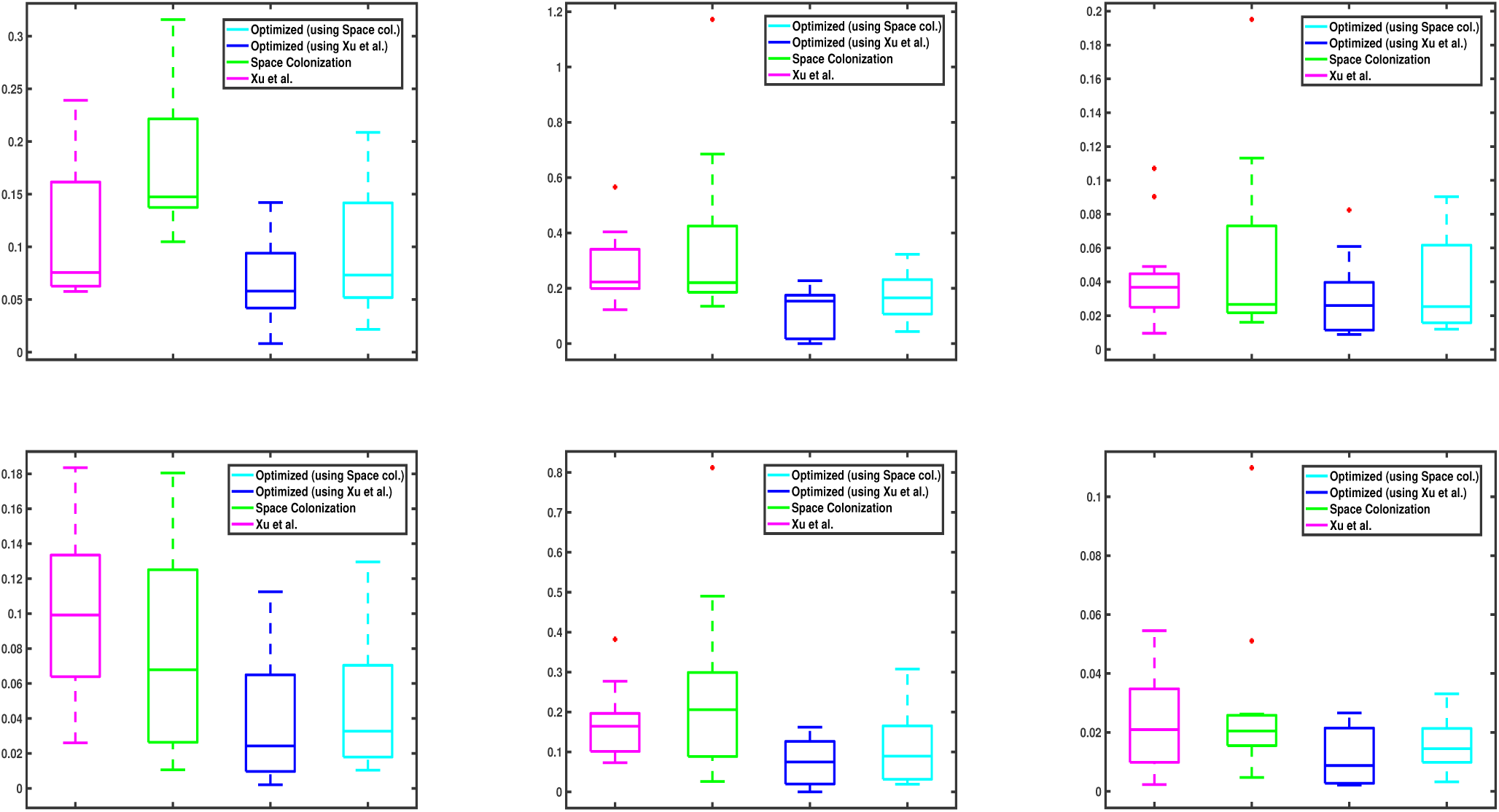
Box plots corresponding to the data values in Figure 5. Top and bottom row shows the relative results of junction point location and branch segment length errors with respect to the groundtruth, respectively. In each graph, the central mark of each box indicates the median, while the bottom and top edges of the box indicate the 25-th and 75-th percentiles of the data, respectively. The whiskers extend to the most extreme data points which are not considered as outliers. The red ‘+’ symbols plot the outliers in the data, if there is any.

## V. Discussion

In this work, we have proposed a technique to improve an existing skeleton by a stochastic optimization technique. The idea of exploiting the original point cloud data in the optimization process allows the skeleton to retain biological relevance with respect to the input point cloud data. The method is tested on different types of data, and improvement over the existing technique is demonstrated. We have demonstrated the effectiveness of the approach by quantifying the junction point location and branch length segment errors, where the branch segment is considered as the length between two consecutive junction points. While the performance varies among the type of the datasets, the optimization clearly improves over the initial skeleton obtained by state-of-the-art ([15], [20], [21]) in the general case. The proposed optimization method can be beneficial to accurate measurements of different types of plant phenotypes.

The method is designed for plants of moderate size having about 100k points. To optimize skeleton of large tree with huge number of points, computation time will be a crucial factor. In these cases, the number of points need to be downsampled to obtain the optimization result in reasonable amount of time. In this work, we did not consider optimizing the computation time or parallelizing the solution for multiple processors. In the current framework, we are not considering the cases of plants with leaves. Although the method might be able to handle the type of leaves having long and narrow shape, but handling other types of complex leaf shapes will be challenging. Given the large variety of leaves of different types of plant species, considering the general case of any type of leaf is beyond the scope of the current work. Another crucial factor is the initial skeleton, which is used as the starting point of the optimization. We have demonstrated that using the method of [15] can be reliably used as the initial skeleton for varieties of cases. However, other types of methods can also be used to compute the primary skeleton, given the fact that the skeleton is reasonably good. As shown in the results, optimization using the Space Colonization ([20], [21]) as the initial skeleton is also performed. However, the results are slightly worse than using the method of Xu *et al*. The reason is that, the optimization result is dependent on the starting point. If the initial skeleton points are too far from the centerline (which results in many cases of Space Colonization algorithm), the optimization fails to move the points to the exact center. Basically the initial skeleton points which are far away, results in getting the optimization to stuck to a bad local minima. In the current formulation we did not consider handling these type of cases, and we assume that the initial skeleton is reasonably correct. The optimization parameters *λ* and *β* play an important role on the final result. By default we keep the values as *λ* = 5, *β* = 5. We tested with different values of *λ* and *β* between the range 1 to 15, and in general the best results are obtained with the default values as stated above. However we notice that for larger trees, higher values of *λ* and *β* slightly improve the result. For any particular species, using a fixed set of values of the parameters should be sufficient.

The centerline assumption of the Gaussian Mixture Model is tested on small plants. However we have not tested the optimization performance for very thick stems, where the skeleton curve might have discontinuous derivatives or high curvature near the branching points. In order to reconstruct the geometry of the branching structure by the notion of generalized cylinders, it will be problematic with the high curvature areas. Handling these types of cases are not considered here, and the framework might need to be reformulated for considering more complex and thicker branching structures. With the goal of reconstructing the geometry of the original plant point cloud data using the skeleton, finding a minimum spanning tree that optimizes the connection between the discrete skeleton points in terms of some biological prior might further improve the skeleton structure. However this is a challenging problem, and we leave it as a future work. Similar type of problem is handled by some heuristic approach in [24], but a generic solution to the problem can be a major part of plant geometry reconstruction.

The proposed method considers plants having no leaves. Skeletonization by considering plant leaves will be worth studying in order to reconstruct the geometry of leafy plants. We have not considered the case where there is lot of noise in the data. Also, explicit handling of occlusion is not modelled in the framework.

## Acknowledgement

This work is supported by Robotics for Microfarms (ROMI) European project. We would like to thank Frédéric Boudon for useful discussions and making the implementation of PlantScan3D library available for public use. We would also like to thank Julie Charlaix for collecting and sharing the data, and other members of the ROMI project for the (not yet published) pipeline to reconstruct the point cloud from images.

## Footnotes

1 https://github.com/fredboudon/plantscan3d

2 By *control points*, we mean the given set of discrete input points, whereas by *spline points* we mean the generated points on the spline curve.

